# Longitudinal dynamics of gut plasmidome and antibiotic resistance during antibiotic therapy: a case report

**DOI:** 10.1101/2025.04.12.648487

**Authors:** Ilmur Jonsdottir, Sean Meaden, Paulina Salminen, Teemu Kallonen, Janne Ravantti, Annaleena Pajander, Sanja Vanhatalo, Matti Jalasvuori, Lotta-Riina Sundberg, Edze Westra, Stineke van Houte, Antti J. Hakanen, Reetta Penttinen

## Abstract

The human gut microbiome is composed of diverse microbes, and its association with human health is well-recognized. Antibiotic therapies for treating infectious diseases often mediate adverse influences on gut microbial ecosystems. Further, ESBL (extended-spectrum beta-lactamase)-producing bacteria potentially residing in the gut benefit from antibiotic-induced environmental changes via positive selection for resistance traits. As antimicrobial resistance (AMR) genes are often harbored by mobile plasmids, the importance of these extrachromosomal mobile genetic elements during antibiotic exposure is evident. However, there is still a knowledge gap in how microbiomes respond to antibiotic treatment especially in terms of plasmid carriage and how gut plasmid populations evolve following and revive after antibiotic therapy. To address these questions, in this case report we investigated the changes in plasmid population in the gut microbiome of a single ESBL-carrying patient during antibiotic therapy for uncomplicated acute appendicitis. Employing longitudinal sampling, we collected *E. coli* strains and performed metagenomic analysis before, during, and after the treatment. Our findings indicate that the antibiotic treatment is associated with a transient alteration in the microbial composition, AMR profile, and plasmid population. An extensive but temporary domination of ESBL-*E. coli* within the gut microbiome was observed parallel to the ongoing antibiotic treatment. However, this was not sustained in the follow-up period, indicating a slight restoration of both the microbial composition and the plasmid population. The research underscores the temporary impact of antibiotic therapy on the dynamics of the gut plasmidome which essentially mediate the spread of AMR within the gut microbiome.

## Introduction

The antimicrobial resistance (AMR) poses a substantial global threat to healthcare^1,2^. In general, the increased emergence of resistant pathogens is due to the widespread usage of antibiotics in healthcare, veterinary medicine, and agriculture. Antibiotic therapy is known to alter and reduce the bacterial diversity of the gut microbiome, as well as to promote resistance by favoring the bacteria carrying resistance traits. Consequently, antibiotics cause disturbance in the gut microbiota, leading to even long-term changes in its composition and functionality^3,4^. This can result in adverse health effects and complications, including colonization by pathogenic bacteria^3,5^. Furthermore, AMR genes are often located in mobile (conjugative) plasmids, which may significantly complicate the treatment of bacterial diseases via disseminating AMR genes within microbial populations^5–10^. However, the epidemiological relevance of AMR plasmids during antibiotic treatment remains unclear^11^. Evaluating the effects of antibiotic usage from an ecological perspective allows to address the role of plasmids in preserving gut microbiota^3,5,10,12,13^.

*Enterobacteriaceae* commonly carry conjugative plasmids that encode AMR genes (ARGs), which can cause serious complications during antibiotic therapy^14–20^. *Escherichia coli,* a member of the *Enterobacteriaceae* family, can be found to both beneficially colonize the human gut as well as to cause a multitude of serious intestinal but also extra-intestinal infections^21^. ESBL-producing *E. coli* strains resist several beta-lactam antibiotics, and therefore they pose a significant clinical challenge. While ESBL-producing bacteria are among the major causes of multiresistant infections, they can also be found harmlessly colonizing the gut microbiome (also referred to as ESBL-carriage). However, in some instances, carriage can lead to limitations on effective antimicrobial therapy, excessive hospital stays, increased healthcare costs, morbidity and mortality ^22–26^. Since antibiotic use exerts selective pressure, promoting the persistence and transfer of ARGs, it becomes imperative to describe the dynamics of the plasmid population during antibiotic therapy. This exploration is pertinent not only in understanding responsiveness to antibiotic therapy but also in assessing post-therapy microbiome recovery, highlighting the broader implications of plasmid dynamics in the context of antibiotic resistance. Furthermore, despite experimental research establishing plasmids as crucial players in antimicrobial resistance and, therefore, in combatting it, our review of the literature revealed a scarcity of longitudinal clinical research describing the impact of antibiotic therapy on plasmid dynamics within the gut microbiota of a human patient^11,27,28^.

Acute appendicitis is one of the most common cause for emergency surgeries worldwide and often associated with pathogenic *E. coli* ^29,30^. The previous randomized controlled trials (RCTs) by the APPAC (APPendicitis ACuta) study group have shown that antibiotic therapy can be a safe and efficient treatment for computed tomography (CT)-diagnosed uncomplicated acute appendicitis ^31–33^. In this study, we used temporal metagenomic analysis to examine the plasmid dynamics and the evolution of antibiotic resistance during antibiotic treatment (ertapenem, levofloxacin and metronidazole) as a case study approach in an ESBL-carrying patient participating in the APPAC III and Microbiology APPAC (MAPPAC) trials (clinical trials done by the APPAC study group) ^34,35^. In addition to studying the impact of antibiotic therapy on the gut plasmid population dynamics, we focused more specifically on *E. coli* bacterial strains isolated during the antibiotic treatment. Our findings indicate that the antibiotic treatment induces a transient alteration in the microbial composition, plasmid population, and AMR profile of the gut microbiome. The antibiotic treatment was followed by an extensive, although temporary, domination of ESBL-*E. coli* (harboring *bla*CTX-M-15 ESBL gene) within the gut microbiome. This was confirmed by a repeated isolation of the same *E. coli* strain before and during the treatment. However, after 180- and 365-day follow-up monitoring, the microbial composition and the plasmid population were partially restored, and the ESBL-*E. coli* (*bla*CTX-M-15) was no longer detected. Instead, other ESBL-*E. coli* (carrying AmpC mutation and beta-lactamases *bla*TEM and *bla*CMY) were observed to colonize the gut microbiome. These results indicate that antibiotic therapy initially leads to a collapse in the diversity of the gut plasmidome, followed by a subsequent recovery, highlighting the dynamic nature of the plasmid population in the microbiome. This study provides a detailed analysis events in a single patient during and after antibiotic treatment, filling a gap with regard to longitudinal plasmid dynamics. These results serve as a starting point for more comprehensive studies aimed at understanding the broader implications of antibiotic treatment on gut microbiota and resistance mechanisms.

## Materials and Methods

### MAPPAC study design and research ethics

Microbiology APPendicitis Acuta (MAPPAC, NCT03257423) prospective cohort study was initiated in 2017 and based at Turku University Hospital with the participation of 10 study hospitals in Finland. The MAPPAC trial enrolled patients concurrently with APPAC III RCT (randomized controlled trial), led by the same study group^36,37^. The MAPPAC protocol corresponds to guidelines from the Standard Protocol Items: Recommendations for Interventional Trials statement^38^. Patients with CT-proven acute appendicitis at Turku University Hospital were informed of all ongoing RCT (APPAC III) and the MAPPAC study. Patients invited to participate in the RCT were also invited to join the MAPPAC trial. Recruited patients were asked to sign a separate consent form for the MAPPAC trial for the usage and collection of their data and microbiological samples. Briefly, APPAC III was a double-blind, placebo-controlled, superiority pilot RCT (NCT03234296) comparing antibiotic therapy (intravenous ertapenem followed by oral levofloxacin and metronidazole) with a placebo to treat uncomplicated appendicitis ^39^. Eligible patients were adults (18 – 60 years of age) with CT-diagnosed uncomplicated appendicitis. The study was approved by the Finnish Medicines Agency and by the Ethics Committee of the Hospital District of Southwest Finland (Turku University Hospital, approval number ATMK:142/1800/2016).

### Patient treatment and sample collection

The patient involved in this case study was a participant in the APPAC III trial and received antibiotic treatment for CT-diagnosed uncomplicated acute appendicitis. The patient treatment was unblinded only after the primary endpoint of the study was completed. The antibiotic treatment lasted seven days: intravenous carbapenem antibiotic ertapenem 1 g once daily for the first three days, followed by four days of fluoroquinolone antibiotics with oral levofloxacin 500 mg once daily and antiprotozoal agent metronidazole 500 mg three times per day. At ten months, additional antibiotic treatment (cephalexin) was orally administered for seven days due to pharyngitis.

A total of six timepoints were sampled from the patient that underwent antibiotic therapy. Rectal swabs were collected at the hospital from the patient at three time points: 1) day 0 (in the emergency department prior to antibiotic treatment), 2) day 1 of treatment, 3) day 3 (change of antibiotics). Fecal samples were collected by the patient at home also at three time points and mailed to the laboratory: 4) day 7 (last day of treatment), and after 5) 180 days and 6) 365 days of the treatment. For timepoints 0, 1, and 3 days, two rectal samples were collected, the first with the Amies-Coal transport swab to allow for bacterial culturing (Sarstedt, Nümbrecht, Germany) and the second with either the Puritan DNA/RNA swab (Puritan Medical products, Guilford, Maine) with shield fluid or with the DNA/RNA Shield Faecal Collection Tube (Zymo Research, Irvine, Canada) for microbial DNA extraction. Shield fluid samples were stored at room temperature for 3-7 days before isolation. The rectal samples were transferred to the laboratory for processing after sampling. For timepoints 7 days, 180 days, and 365 days, the patient collected fecal samples in a 10 mL DNA/RNA Shield Faecal Collection Tube (Zymo Research), enabling transportation and storage at room temperature before DNA extraction. A second sample was also collected in a gel tube during these timepoints for culture purposes.

### *E. coli* sampling, strain isolation, and characterization

Bacterial samples from Amies-Coal swabs were cultured on CHROMagar™ Orientation (Becton Dickinson, Heidelberg, Germany) plates from where *Escherichia coli* resembling pink colonies were picked for pure culture. Pure cultures were also performed on Chromagar orientation plates, and bacterial strains were stored at -80 °C.

Bacterial cultures were grown in Luria-Bertani media at 37°C (with 200 rpm agitation for liquid cultures) ^40,41^. The isolated *E. coli* strains were screened for ESBL by streaking on chromogenic ESBL plates (ChromID ESBL, Biomerieux). Antimicrobial resistance and susceptibility tests were performed using agar plates supplemented with lethal doses of appropriate antibiotics (50 µg/ml cephalothin, 150 μg/ml rifampicin, 150 μg/ml ampicillin). Mobility of ESBL gene was performed through an overnight coculture of the rifampicin-susceptible ESBL donor strains and three rifampicin-resistant *E. coli* strains (HMS174 (Navagen, Madison, WI, USA), DSM4860 (DSMZ, Braunschweig-Süd, Germany), and DSM8696 (DSMZ, Braunschweig-Süd, Germany)) (see Supplementary Table S1). ESBL-transconjugants were selected on agar plates supplemented with 150 μg/ml rifampicin and 50 μg/mL cephalothin.

### Isolate sequencing and bioinformatic analyses

DNA from all the bacterial isolates was extracted using the Wizard Genomic DNA purification kit (Promega) according to the manufacturer’s instructions. The DNA concentration was determined with a Qubit^TM^ 3.0 fluorometer using the dsDNA HS kit (InvitrogenTM, ThermoFisher Scientific). The sequencing libraries were prepared using a Short-insert library and sequenced with the DNBSEQ platform (PE150). The reads were initially quality- and adapter-trimmed with SOAPnuke by the sequencing service; reads containing more than 1% of N, more than 40% of the bases in a read have quality value under 20, or reads with length under 150 bp were removed^42^. The reads were further trimmed with Trimmomatic (v0.39), and reads with lengths under 100 bp after trimming were discarded from the analysis^43^. Quality control was performed through FastQC (v 0.12.0)^44^. The trimmed reads were *de novo* assembled using SPAdes (v 3.15.5)^45^. The resulting sequences were annotated with Prokka rapid prokaryotic genome annotation (v 1.4.6)^46^. CRISPR defense mechanisms were identified from all strains using the web server CRISPRCasFinder and PADLOC (v 1.1.0)^47,48^.

Resistance genes and mutations were discovered with Resfinder 4.1 (Centre for Genomic Epidemiology)^49–51^. Plasmid sequences were identified and extracted from the assembled contigs using geNomad^52,53^. The plasmid sequences were clustered based on 95% average nucleotide identity (ANI) and 85% alignment fraction (AF) of the shorter sequence following the approach of Nayfach et al. (2021)^54^, which uses blast+ to compute an all-vs-all local alignment then processes the resulting alignments to identify the shared plasmids among strains. The representative sequence of each cluster was analyzed with COPLA plasmid taxonomic classifier tool^55^ to identify plasmid incompatibility (Inc) type, plasmid taxonomy unit (PTU), closest plasmid reference sequence based on ANI (average nucleotide identity) and BLASTN search^51,56^. Plasmid sequences were considered as true plasmids if they 1) carried at least one plasmid marker gene (assessed by geNomad), and 2) were identified to carry an Inc replicon site or classified into PTU (by COPLA). The sequences resulting in the same BLAST hit results were considered to originate from same plasmid and treated as one. The *E. coli* sequence type was determined through MLST 2.0 (Centre for Genomic Epidemiology) ^51,57–62^. Phylogenetic tree was inferred by CSI Phylogeny 1.4 (Centre for Genomic Epidemiology)^63^ with a reference ST648 genome (GenBank: CP008697.1). The plasmid carriage was visualized with the phylogenetic tree using iTOL (Interactive Tree of Life)^64,65^.

### Metagenomic sequencing and bioinformatic analyses

Whole Genome Shotgun sequencing was performed on the DNA extracted with a semiautomated GXT Stool Extraction kit (Hain Lifescience GmbH, Nehren, Germany) from the DNA/RNA Shield Collection Tube w/ Swab (Zymo Research) and DNA/RNA Shield Fecal Collection Tube (Zymo Research). Metagenomic shotgun sequencing of the gut microbiota was performed using Nextera XT DNA Library Preparation Kit (Illumina) from a total DNA input of 1 ng and sequenced on an Illumina HiSeq X platform 150PE run at CosmosID (Maryland, USA). The reads were trimmed with fastp^66^. Host read removal was performed by mapping the reads against the reference human genome with bowtie2 and utilizing only the unmapped reads for further analysis ^67^. For any quantitative analysis, the reads were subsampled based on the lowest read number (5 553 035 of paired reads at sampling point 3) with seqtk^68^. The microbial community composition was determined from the subsampled reads with kraken2 using the standard database after removing reads classified as human^69,70^. Phylum-level classifications were extracted and used to generate Shannon’s alpha diversity index and Bray-Curtis distances using the ‘vegan’ package, and the ‘ecodist’ package was used for principle coordinate analysis in R (version 4.3.0)^71,72^. Rarefaction curves were generated for the total data from each sample using the ‘rarecurve’ function in ‘vegan’ (Fig. S1). To assess which taxa were driving differences between the samples, non-metric multidimensional scaling (NMDS) analysis was used with the ‘envfit’ function in ‘vegan’ to plot the species loadings (Fig. S2).

All the metagenomic reads from all samples were co-assembled with metaSPAdes (v 3.15.5) ^73^, and the assembled contig coverage was assigned with bowtie2 and CoverM ^67,74^. The contigs were binned with metaBAT to produce metagenome-assembled genomes (MAGs)^75^. The MAG completeness was confirmed by checkM^76^. The plasmid contigs were identified from the co-assembled contigs with geNomad^52,77^. Plasmid sequences were clustered using the same methods as the isolate plasmid sequences (95% ANI, 85% AF) following the protocol described previously^54^. Subsampled reads were mapped to these clustered plasmid sequences, coverage measured with CoverM^74,78^ using the ‘metabat’ metric, and the resulting data converted to a plasmid abundance table for downstream analysis. The diversity analyses of the plasmid populations were analyzed using the same methods as the whole community data, again using the ‘vegan’ and ‘ecodist’ R packages. For plasmid dynamics visualization, the plasmid contigs were filtered by excluding any contig shorter than 750 bp, with a threshold of at least one plasmid marker gene (assessed by geNomad) and minimum average coverage of 1 in at least one of the samples and visualized with the ‘ggalluvial’ R package^79^. In order to identify *E. coli* isolate plasmids from the metagenomic data, all plasmid sequences (from isolates and metagenomic data) were clustered together using the same methods as above (95% ANI, 85% AF). Resistance genes were subsequently identified from the co-assembled contigs using ResFinder 4.1.

## Results

### Characteristics of *E. coli* strains isolated during antibiotic treatment

We investigated the impact of antibiotic therapy on the gut plasmidome of a single patient via longitudinal sampling during a 7-day ertapenem-levofloxacin-metronidazole treatment. The samples were collected as part of the MAPPAC study at each time point: day 0, day 1, day 3, day 7, day 180 and day 365. The samples were utilized for the isolation of *Escherichia coli* strains and metagenomic shotgun sequencing (excluding day-1 sample) to determine the dynamics of bacterial, plasmid, and AMR populations at the community level.

Twelve *E. coli* strains were isolated across six timepoints. These *E. coli* isolates were subjected to phenotypic resistance and plasmid mobility testing, as well as whole genome analysis, encompassing Multi-Locus Sequence Typing (MLST) -based sequence typing, antimicrobial resistance profiling, plasmid carriage assessment, and analysis of CRISPR-Cas systems (see Table 1). The same strains of *E. coli* were consistently isolated throughout the antibiotic therapy from day 1 to 7, persisting even after the antibiotic cycling on day 3. All *E. coli* isolates underwent genotypic profiling using MLST, categorizing them into five distinct Sequence Types (STs). The *E. coli* strains isolated before and during treatment exhibited exclusively two ST types: ST648, carrying the *bla*CTX-M-15 gene, and a non-resistant ST6467 strain. In contrast, the isolates obtained from the subsequent follow-up assessments at 180 days and 365 days had not been previously observed at earlier time points. They represented three distinct STs: ST929 (n=1), ST69 (n=2), and ST73 (n=1). It is noteworthy that specific STs, such as ST69 and ST73, are recognized as established pandemic lineages within the category of extra-intestinal pathogenic *Escherichia coli* (ExPEC)^80–85^

**Table 1.** *E. coli* strains isolated during antibiotic treatment. Genetic features related to antimicrobial resistance (AMR including antibiotic resistance genes (ARGs), plasmid carriage and CRISPR-Cas system.

| Isolate | Timepoint | MLST | AMR |  |  |  | Plasmid(s) | CRIS PR type |
| --- | --- | --- | --- | --- | --- | --- | --- | --- |
|  |  |  | ARGs |  | Mutations |  |  |  |
|  |  |  | Beta-lactamase | Other | ESBL | Other |  |  |
| 0.1 | Before treatment | 6467 | None | None | None | *gyrA*, *gyrB*, *parC*, *pmrA*, *folP*, *rpoB*, 16S-*rrsH* | None | IF |
| 0.2 | Before treatment | 648 | *bla*CTX-M-15 | *sul2* | AmpC-promoter | *gyrA*, *gyrB*, *parC*, *parE*, *pmrA*, *pmrB*, *folP*, *rpoB*, 16S-*rrsB/C/H* | Col(BS512), IncFII, IncI2(Delta) | IE |
| 1.1 | Treatment (day 1) | 648 | *bla*CTX-M-15 | *sul2* | AmpC-promoter | *gyrA*, *gyrB*, *parC*, *parE*, *pmrA*, *pmrB*, *folP*, *rpoB*, 16S-*rrsB/C/H* | Col(BS512), IncFII, IncI2(Delta) | IE |
| 1.2 | Treatment (day 1) | 6467 | None | None | None | *gyrA*, *gyrB*, *parC*, *pmrA*, *folP*, *rpoB* | None | IF |
| 3.1 | Treatment (day 3) | 648 | *bla*CTX-M-15 | *sul2* | AmpC-promoter | *gyrA*, *gyrB*, *parC*, *parE*, *pmrA*, *pmrB*, *folP*, *rpoB*, 16S-*rrsB/C/H*, 23S | Col(BS512), IncFII, IncI2(Delta) | IE |
| 3.2 | Treatment (day 3) | 648 | *bla*CTX-M-15 | *sul2* | AmpC-promoter | *gyrA*, *gyrB*, *parC*, *parE*, *pmrA*, *pmrB*, *folP*, *rpoB*, 16S-*rrsB/C/H*, 23S | Col(BS512), IncFII, IncI2(Delta) | IE |
| 7.1 | Treatment (day 7) | 648 | *bla*CTX-M-15 | *sul2* | AmpC-promoter | *gyrA*, *gyrB*, *parC*, *parE*, *pmrA*, *pmrB*, *folP*, *rpoB*, 16S-*rrsB/C/H* | Col(BS512), IncFII, IncI2(Delta), Col440I | IE |
| 7.2 | Treatment (day 7) | 648 | *bla*CTX-M-15 | *sul2* | AmpC-promoter | *gyrA*, *gyrB*, *parC*, *parE*, *pmrA*, *pmrB*, *folP*, *rpoB*, 16S-*rrsB/C/H* | Col(BS512), IncFII, IncI2(Delta) | IE |
| 6.1 | 6-month follow-up (180 days) | 929 | None | *sit*ABCD | AmpC-promoter | *gyrA*, *gyrB*, *parC*, *parE*, *pmrA*, *pmrB*, *folP*, *rpoB*, 16S-*rrsB/C/H*, 23S | Col440I | None |
| 6.2 | 6-month follow-up (180 days) | 69 | *bla*TEM-1B*, *bla*CMY-variant* | *qnrS1*, *mph*(A), *sul2*, *dfrA1*, *tet*(A), *aadA1*, *sit*ABCD, *aph*(6)-Id, *aadA1*, *aph*(3")-Ib, *qacE* | AmpC-promoter | *gyrA*, *gyrB*, *parC*, *parE*, *pmrA*, *pmrB*, *folP*, *rpoB*, 16S-*rrsB/C/H*, 23S | IncB/O/K/Z, IncFIB, IncFII, IncX4 | IE |
| 12.1 | 1-year follow-up (365 days) | 69 | *bla*CMY-variant* | *sul2*, *sit*ABCD, *aph*(6)-Id, *aph*(3")-Ib | AmpC-promoter | *gyrA*, *gyrB*, *parC*, *parE*, *pmrA*, *pmrB*, *folP*, *rpoB*, 16S-*rrsB/C/H*, 23S | IncB/O/K/Z, IncX4 | IE |
| 12.2 | 1-year follow-up (365 days) | 73 | None | *sit*ABCD | AmpC-promoter | *gyrA*, *gyrB*, *parC*, *parE*, *pmrA*, *pmrB*, *folP*, *rpoB*, 16S-*rrsB/C/H*, 23S | Col156, IncFIB, IncFII, IncFII(29) | None |

\* Located on a plasmid

In our investigation of ESBL occurrence among these *E. coli* isolates, genetic and phenotypic testing revealed that 9 of the 12 isolates (67%) were found with ESBL-producing characteristics (Table 1, Supplementary Table S1). None of the isolates showed ESBL mobility with the tested recipient strains (Supplementary Table S1). Notably, 10 of the strains (representing sequence types ST69, ST73 ST648 and ST929) carried mutation(s) in AmpC promoter, which is often associated with ESBL properties as a result of overexpressed *ampC* gene ^86,87^. However, regardless of the AmpC promoter mutations, strain 6.1 (ST929) did not exhibit ESBL resistance, but was susceptible to ampicillin and cephalothin (Supplementary Table S1). ESBL gene *bla*CTX-M-15 was identified in six of the isolates which also shared commonalities in terms of resistance genes, sequence type, plasmid profile, and CRISPR-Cas system. Further, the phylogenetic analysis based on whole genome SNPs (Figure 1) showed that all six ST648 isolates, identified at timepoints 0, 1, 3, and 7 days, are likely to be isolates of the same strain. This led us to infer that the same strain persisted throughout antibiotic therapy. Sulfonamide resistance gene *sul*2 was consistently detected at all time points, in 67% (n=8) of the isolates. One particular *E. coli* isolate, denoted as 6.2, displayed the highest resistance profile among the strains and was obtained during the 180-day follow-up assessment. Notably, this isolate (6.2) bore 13 distinct resistance determinants, including two beta-lactamase genes (*bla*TEM-1B and a *bla*CMY-2 variant), the *sul*2 gene, *tet*(A), conferring tetracycline resistance, and *qnr*S1, imparting low-level resistance to quinolones.

**Figure 1.**
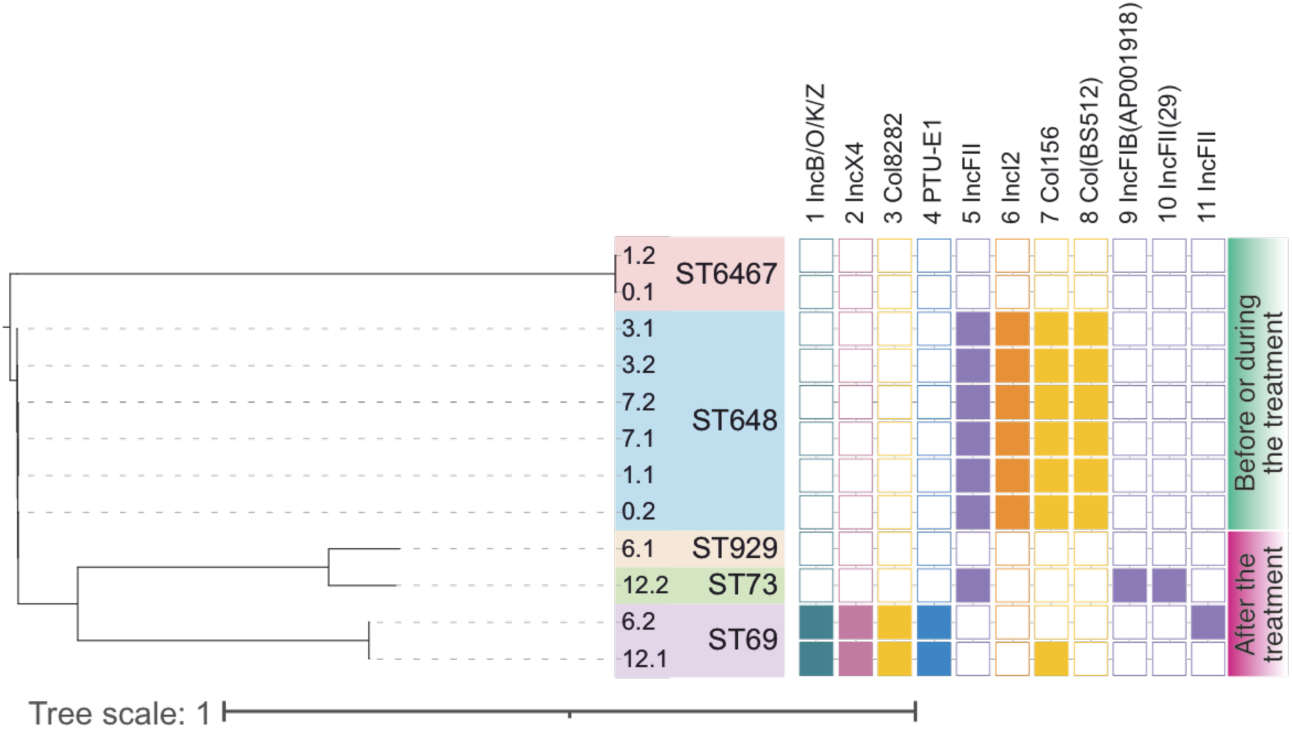
Plasmid carriage of the isolated *E. coli* strains and their sequence types (ST). Phylogenetic tree is inferred from whole genome SNP (CSI Phylogeny). Plasmids were identified with geNomad, clustered within all strains and filtered with the following criteria: >1 marker gene and either/or Inc type detected and PTU (plasmid taxonomic unit) assigned. Plasmid identifier (1-11) refers to plasmid clusters, for more detailed information on plasmid characteristics, see Supplementary Table S2. Plasmid Inc type is shown, except for plasmid 4 where Inc type was not identified and PTU is shown instead. Plasmid carriage per each plasmid cluster is shown as color-filled square.

We further explored the plasmid burden of the *E. coli* isolates. After plasmid sequence classification, clustering and filtering, a total of 72 plasmid clusters originating from at least 11 plasmids (Figure 1, Supplementary Table S2) were identified. One of the plasmids, plasmid identifier #1, was detected as IncB/O/K/Z plasmid and was observed to harbor several antimicrobial resistance genes (betalactamase *bla*CMY, aminoglycoside resistance genes *aph(6)-Id* and *aph(3’’)-Ib*, and *sul2* sulfonamide resistance gene). Plasmid #1 was exclusively detected in the follow-up timepoints at 180 and 365 days. Notably, all ST648 isolates (0.2, 1.1, 3.1, 3.2, 7.1, and 7.2) shared the same plasmid profile and each of these strains were isolated before or during the treatment. Conversely, no plasmids were detected in ST6467 (isolates 0.1 and 1.2), and correspondingly, these isolates did not exhibit any resistance traits, and based on phylogenetic analysis are isolates of the same strain (Figure 1). From ST929 (strain 6.1) no contigs assigned as plasmids were found, however plasmid marker for Col440 was identified in the isolate’s whole genome analysis. Furthermore, the phylogenetic analysis of isolates 6.2 and 12.1 (ST69) suggested they were the same or closely related strain, exhibiting similar but not identical plasmid profiles. The difference found was in the plasmid-encoded *bla*TEM-1B and *qnr*S1 genes, unique to isolate 6.2. Isolates 6.1 (ST929) and 12.2 (ST73) both possessed plasmids. However, their plasmid profiles differed from those observed in the other isolates.

### Effect of antibiotic treatment on gut microbial community composition

The collected fecal samples (day 0, day 3, day 7, day 180, day 365) were subjected to metagenomic shotgun sequencing analysis (excluding day 1 sample) to determine the dynamics of bacterial and plasmid populations as well as antibiotic resistance profile at the community level. The taxonomic profiling of the metagenomic dataset exhibited dynamic development of the microbial community composition in parallel to the 7-day antibiotic therapy (Figure 2, Supplementary Figures S1-S2). A marked shift in the microbial composition was observed by days 3 and 7 when the antibiotic therapy was ongoing. Interestingly, an increase in microbial alpha diversity was observed on day 3 (Figure 3A). However, by the end of antibiotic treatment on day 7, the microbial community was heavily dominated by the genus *Escherichia*, accounting for over half of the microbial composition (Figure 2B). The diversity of the community also decreased significantly on day 7 (Figure 3A). However, by the follow-up timepoints on days 180 and 365, the *Escherichia* dominance was cleared, and partial recovery of microbial diversity was observed. When we assessed the similarity of communities over time as beta diversity, it was revealed that the community composition shifted on days 3 and 7, respectively with the ongoing antibiotic treatment (Figure 3B). In other words, while the alpha diversity was shown to recover by partial replacement of the original community, the beta diversity analysis clarified that the community structure remained distinct after the 365-day follow-up.

**Figure 2.**
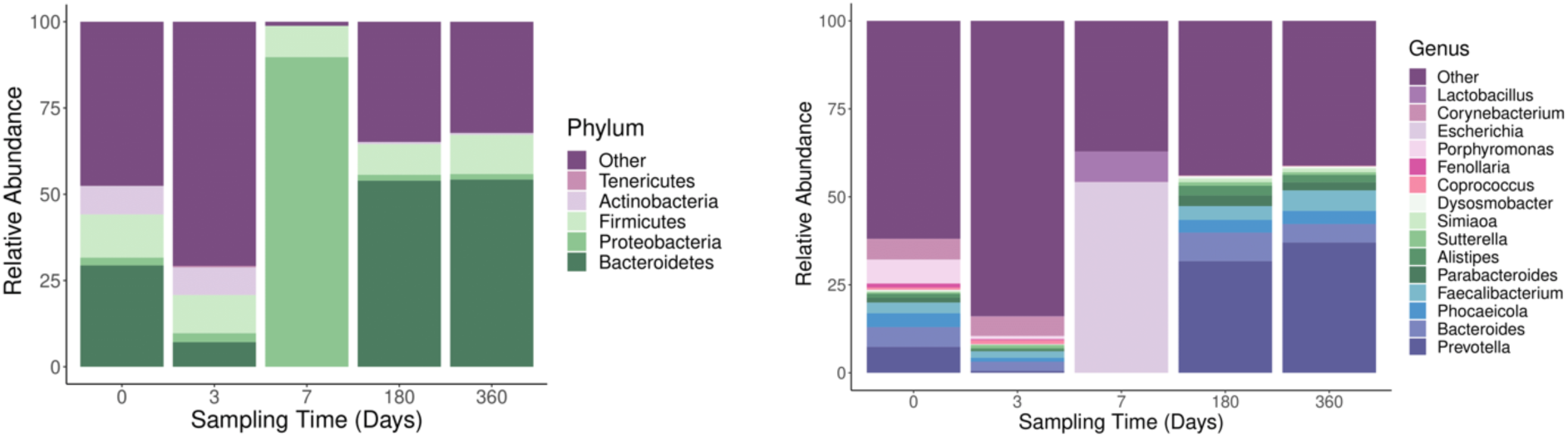
Composition of the microbial community at a phylum (A) or genus level (B). The taxonomy was assigned with kraken2, which classifies reads using the standard database.

**Figure 3.**
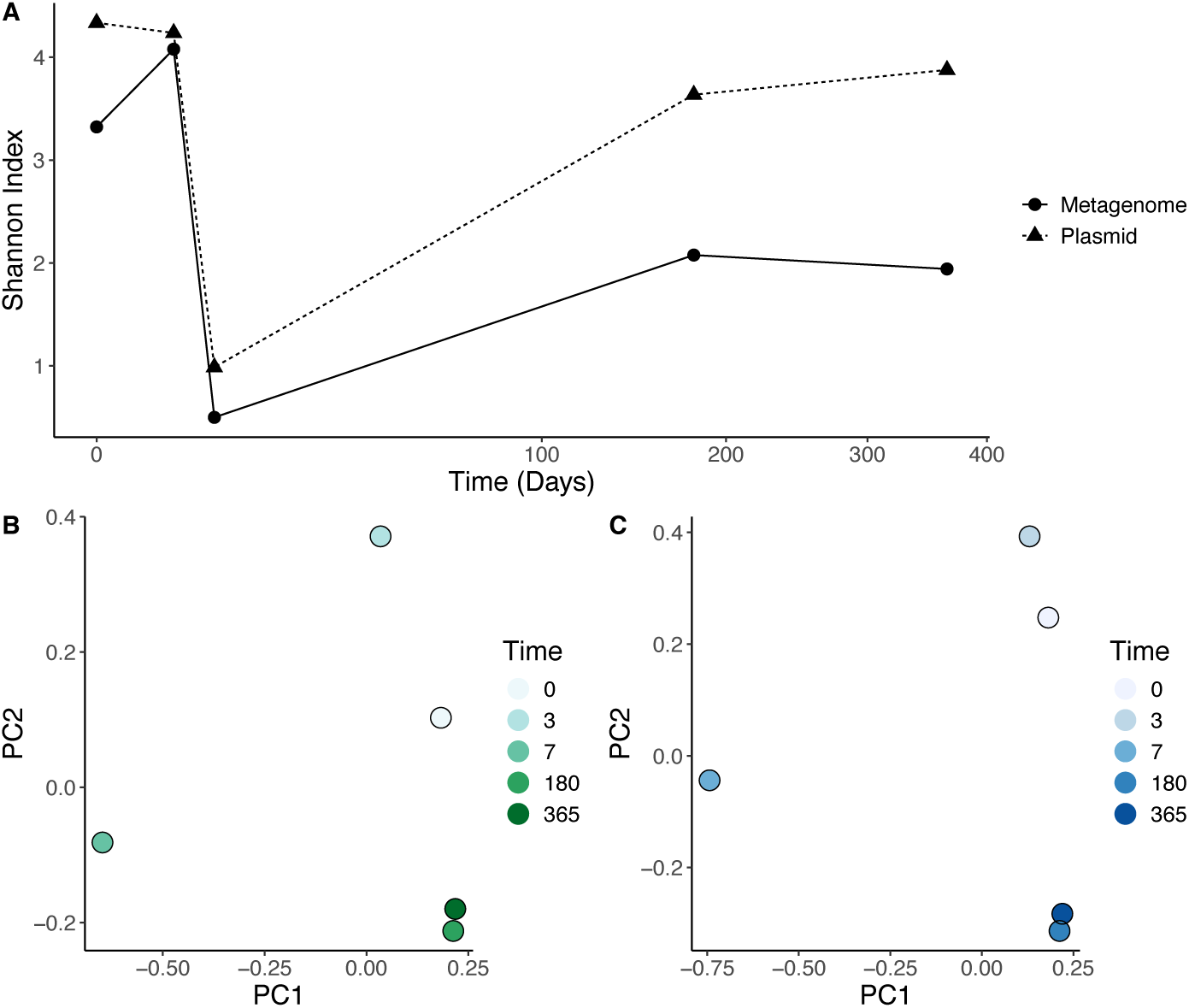
Microbial and plasmid diversity and similarity in the gut microbiome during a 7-day antibiotic treatment determined before (0) and (3, 7, 180, 365 days) after the treatment initiation. The alpha diversity of microbial (at genus level) and plasmid population during the sampling period (A). The recovery of the microbial (B) and plasmid (C) populations after the antibiotic treatment was measured as community similarity by principal coordinate analysis.

### Plasmid dynamics during antibiotic treatment

To elucidate the effects of antibiotic treatment on the gut plasmidome dynamics, we extracted the plasmid contigs from metagenomic assemblies and studied their prevalence over time. A total of 2212 plasmid contigs were identified across the samples after clustering and 335 after filtering based on length and marker gene presence. In general, similar effects were seen in the plasmid population as in the microbial community during antibiotic treatment. A remarkable collapse in plasmid diversity was observed during the antibiotic therapy on days 3 and 7 (Figure 3A). On day 7, the plasmid population was essentially dominated by only a few plasmids (Figure 4). According to our clustering analysis, all of these plasmids were identified to be the same plasmids found from the *E. coli* ST648 that was isolated at several time points before and during the antibiotic treatment. Yet, the plasmid contigs that peaked during the treatment (higher prevalence on days 3 and 7) did not remain dominant within the microbial community after the treatment. Instead, at least partial recovery of the plasmid population by the 180 and 365-day follow-ups was observed. Comparably to the microbial hosts, the plasmid population similarity also shifted during antibiotic treatment. Regardless of the partial recovery in diversity, the plasmid populations were clearly different from the original (before the treatment), as revealed by similarity analysis assessed via beta diversity (Figure 3C). Yet, the plasmid population was very similar after 180 and 365 days (Figure 3C, Figure 4).

**Figure 4.**
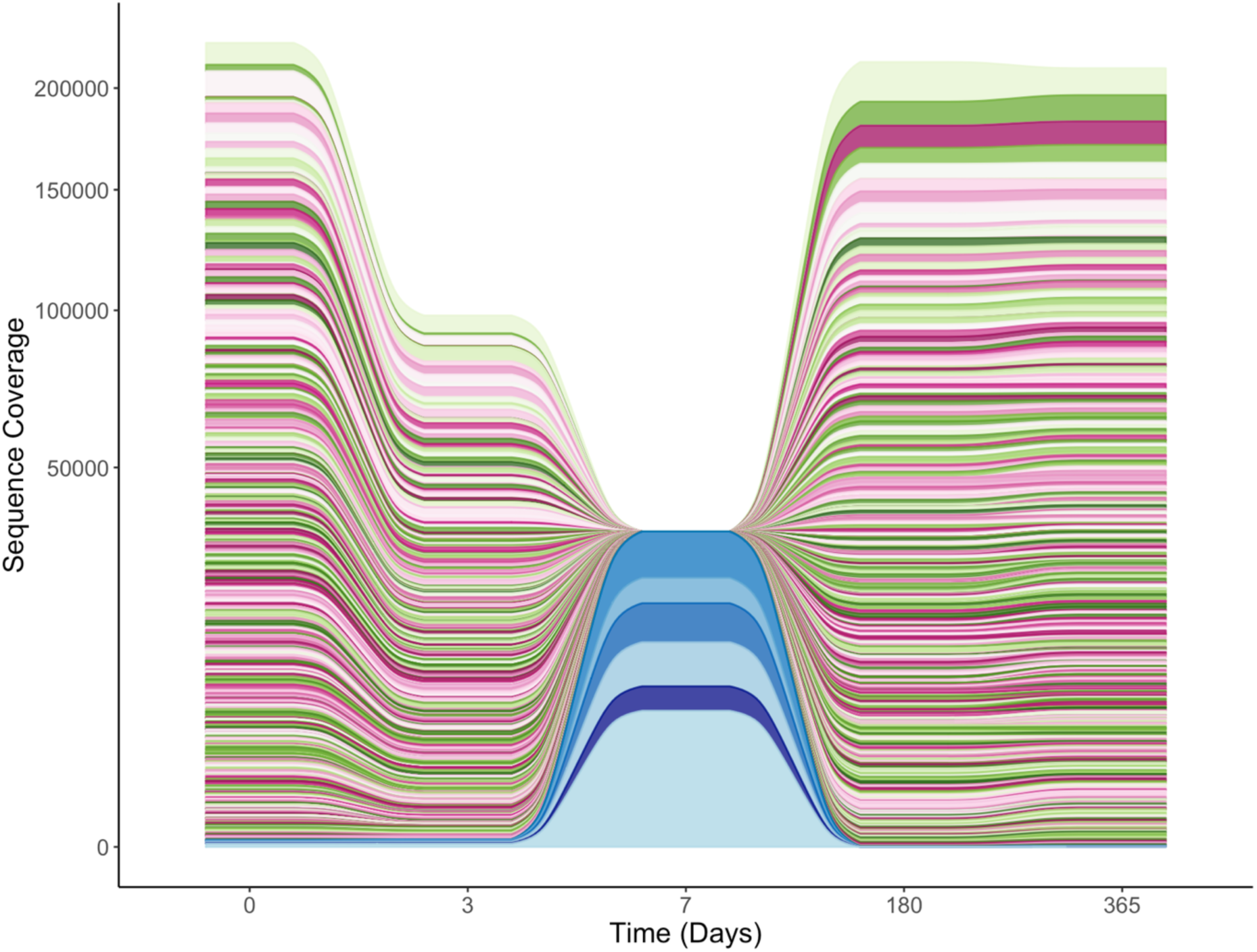
Dynamics of the plasmid population in the patient gut community during a 7-day antibiotic treatment presented as plasmid contig coverage. Plasmids were classified with geNomad, and filtered with minimum length of 750 bp and carriage of >1 marker gene to simplify the visualization. Plasmids found from *E. coli* isolates are shown in blue. Ribbon heights represent sequencing coverage per plasmid at each time point.

We further investigated the plasmid population based on their first occurrence to gain deeper insight into the plasmid recovery but also colonization of new plasmids after antibiotic treatment. Predominantly, the plasmid population after 365 days consisted of focal plasmids (those present in the microbial community before the antibiotic treatment) (Figure 5). However, only a few new plasmids were seen to colonize the microbiome subsequent to the start of antibiotic therapy, as seen on day 3, whereas no new plasmid colonizations were detected on day 7. During the post-treatment surveillance, new colonizing plasmids were mainly recorded on the 180-day follow-up, and some were documented after one year.

**Figure 5.**
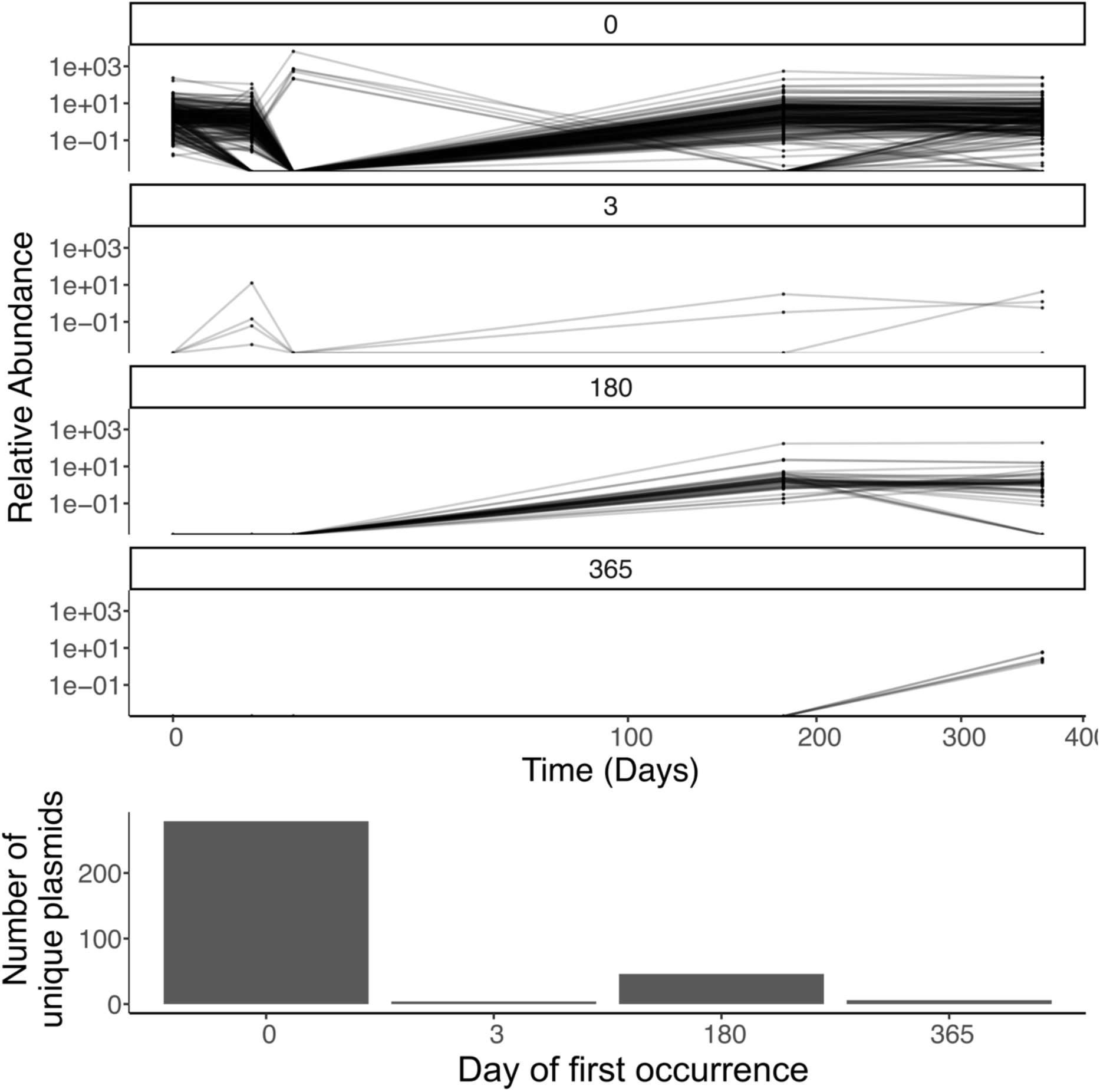
Plasmid abundance based on first occurrence over time. The top panel displays the relative abundance of plasmids at day 0, day 3, 180, and 365 days post-treatment. The bottom panel shows the number of new unique plasmids identified at each time point. Note: No new plasmids were detected at day 7, so this time point is omitted from the analysis.

### Evolution of gut AMR profile

To track the changes in the antimicrobial resistance profile during antibiotic therapy, we extracted the AMR gene-carrying contigs from the metagenomic assemblies, identified their genetic context (plasmid or other), and measured their relative abundance across the time points. The AMR profile exhibited 25 ARGs altogether in 23 contigs, of which 9 were associated with plasmid and 14 with a chromosomal contig (Figure 6). Prominently, two genes (also detected from certain *E. coli* isolates, see Table 1), ESBL gene *bla*CTX-M-15 and sulfonamide resistance gene *sul*2, were barely detectable in the metagenomic data before the antibiotic treatment but were found dominating the gut resistome on day 7. Notably, the genomic locations of these genes varied, with *bla*CTX-M-15 residing on the chromosome, while *sul*2 was located on a plasmid, consistent with the observations from *E. coli* whole genome analysis. Intriguingly, both *E. coli* resistance genes disappeared by the follow-up timepoints after 180 and 365 days.

**Figure 6.**
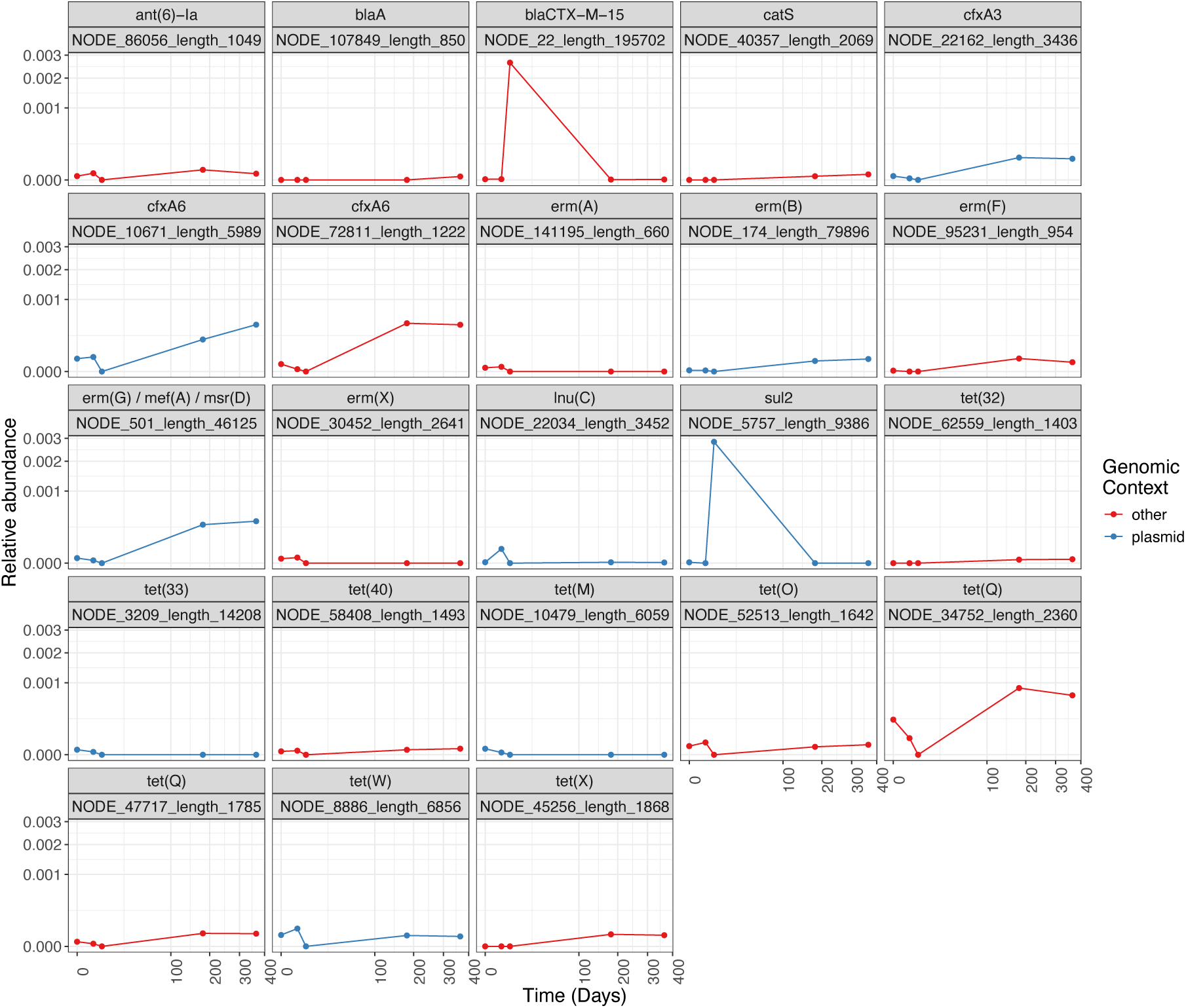
Prevalence of antibiotic resistance genes during antibiotic therapy. Each plot depicts the relative abundance of resistance gene(s) found in a unique location (contig/node), with the contig/node identifier labeled below the name of the resistance gene(s). The genetic context of the resistance genes is distinguished by color: blue for plasmid-associated genes and red for genes located in other contexts (e.g. chromosomal).

Three plasmid-carried macrolide resistance genes, *erm*G, *mef*A, and *msr*D, were localized on the same plasmid contig. This plasmid manifested a modest reduction in abundance after initiating antibiotic therapy, followed by a significant upsurge in the subsequent follow-up time points. Furthermore, the beta-lactamase gene *cfx*A6 was detected in two distinct genetic locations, one putatively residing in a chromosome and the other on a plasmid. Both *cfx*A6 genes demonstrated a substantial increase in abundance following antibiotic treatment at timepoints 180 and 365 days. A comparable increase in abundance was observed for the chromosomally-predicted *cfx*A3 gene.

In our analysis, we identified tetracycline resistance genes in nine contigs, the majority of which being chromosomally located, while three were situated on plasmids, specifically *tet*33, *tet*M and *tet*W. The *tet* gene abundance consistently exhibited varying patterns across the time points, with *tet*32, *tet*33, *tet*40, and *tet*M maintaining relatively low abundance. Conversely, the relative abundance of *tetQ and tetX* genes remained higher in the post-treatment follow-up period compared to the level before antibiotics. Taken together, an evident shift in the resistance profile after the antibiotic treatment was prominently characterized by the increased prevalence of *cfx*A3, *cfx*A6, *erm*G, *mef*A, *msr*D, and *tet*Q resistance genes.

In order to identify the host bacteria of the most prevalent resistance genes after the treatment, we binned the metagenomic contigs to produce metagenomic assembled genomes (MAGs). However, the chromosomally located beta-lactamase gene *cfx*A6 or tetracycline resistance gene *tet*Q were not associated with any of the MAGs and therefore a nucleotide blast (blastn) search was performed to track their potential host species. The contig harboring *cfx*A6 was mainly associated with *Segatella copri* (previously known as *Prevotella copri*) and *tet*Q-harnessing contig with *Bacteroides fragilis*.

The host of the plasmid-associated AMR genes were identified by blastn searching the plasmid contig sequences. All of the AMR plasmid contigs indicated CFB (Cytophaga-Flavobacterium-Bacteroides) group as their potential host bacteria. The 3436 bp contig containing beta-lactamase gene *cfx*A3 was pointed at *Bacteroides* and *Parabacteroides* genomes, however the *cfx*A3 was accompanied by *Parabacteroides* mobilization protein (Genbank: UVQ46764.1). Beta-lactamase gene *cfx*A6 was found from a 5989 bp contig identified as *Bacteroides fragilis* plasmid (GenBank: KJ830768.1) which also carried mobilization protein encoding genes *mobA* and *mobB*. Finally, the 46125-bp plasmid contig identified with several AMR genes (*erm*G, *mef*A, *msr*D) was found to carry several conjugative transposon genes (*traK, traM, traN*) and a type IV secretory system conjugative DNA transfer family protein gene. However, the most closely related hits were found from chromosomes of *Alistipes* and *Bacteroides* species, which suggests that this mobile genetic element may also exist chromosomally integrated in our dataset.

## Discussion

Antibiotics likely lead to significant shifts in community diversity and the antibiotic resistance profile in the gut, which may, in turn, have adverse effects on human health. We examined the outcomes of an antibiotic treatment of an ESBL-carrying patient diagnosed for uncomplicated acute appendicitis. Our analysis employed a combination of metagenomic analysis and the examination of gut *E. coli* strains isolated from corresponding sampling points. In this case study, the metagenomic sequence analysis revealed an apparent decline in the microbial and plasmid diversity between days 3 and 7, putatively reflecting the effect of the 7-day combination antibiotic treatment with ertapenem, levofloxacin and metronidazole. Further, a transient reduction in microbial diversity and the simultaneous enrichment of ESBL-producing *E. coli* was observed by day 7. Levofloxacin is a fluoroquinolone antibiotic with a broad spectrum of activity against both Gram-positive and Gram-negative bacteria^88^. Previously, levofloxacin usage has been reported to provoke fluoroquinolone resistance in *E. coli*^89^. Carbapenems have been reported for their adverse effect on the gut microbiota of infirmed patients, and there is an increasing trend of carbapenem-resistant *Enterobacteriaceae* within hospitalized patients ^90–92^. This seems to correlate with a global rise in carbapenem consumption due to the “pandemic” of ESBL-producing *Enterobacteriaceae*^93^. However, in our study, no known carpapenemase encoding resistance genes were found in the isolated *E. coli* strains nor in the metagenomic data. During subsequent follow-up periods at days 180 and 365, our analysis revealed a partial recovery in microbial and plasmid population diversities. A comparison between 180 and 365 days found notable similarities both in microbiome and plasmidome. Understanding such recoveries holds significance in assessing the risks associated with antibiotic therapy for appendicitis, given the prominent role of the gut microbial community in the pathogenesis of various diseases ^94–98^. The unique role of the appendix adds another layer of complexity. Indeed, the appendiceal microbiota can differ significantly among various patient groups, influencing the diversity and density of the microbial taxa ^99^. Specifically, appendicitis can lead to dysbiosis, characterized by a reduction in microbial diversity and richness, resulting from infection and inflammation^100,101^. Vanhatalo et al. showed that patients with uncomplicated appendicitis exhibited distinct microbial profiles compared to those with complicated cases, illustrating how the condition itself can alter the microbiome composition^99^.

Extensive sequencing and experimental research have unequivocally demonstrated the role played by conjugative plasmids in the dissemination of antimicrobial resistance genes among bacterial populations ^102–105^. Antibiotic pressure has been shown to promote the survival of certain plasmids that confer selective advantages under treatment conditions^106,107^. In our analysis, we observed a substantial decline in the plasmid diversity by the end of antibiotic treatment, mirroring a similar pattern seen in the bacterial population. Interestingly, the few plasmid contigs present on day 7 were identified to be the plasmids carried by the isolated *E. coli* strains. However, while dominant after the treatment, none of these plasmids were present in the metagenome during the follow-up timepoints.

Given the well-known unbalancing effects of antibiotics on gut microbiota, it is acknowledged that a disrupted microbial community is more prone to an invasion by e.g. bacterial pathogens^110,111^. Similarly, the dynamics of plasmid retention and loss in response to antibiotic treatment is well-documented; a sudden reduction in microbial diversity can create niches that may be more readily colonizable by resistant strains, including those carrying novel plasmids^108,109^. This phenomenon is compounded by the possibility for opportunistic acquisition of novel mobile genetic elements, such as plasmids, which may confer additional resistance traits to resident bacterial populations^112^. We examined the plasmidome based on the first occurrence of each plasmid contig to track the colonization events of new invading plasmids. While the decline in plasmid diversity at the end of antibiotic treatment seemed to partially recover, several new colonizing plasmids were observed to persist by the 365 days follow-up.

The emergence of new plasmids in the system may reflect their ability to capitalize on the transient gaps in the microbial population created by antibiotic disturbance. A growing body of research underscores that antibiotic treatment not only eradicates sensitive strains but can also facilitate the uptake of foreign resistance plasmids as a response of altered surviving bacterial communities harboring different plasmid repositories^113,114^. The analysis for plasmid beta diversity aligned with this assumption, as 180 and 365-day plasmid populations clustered apart from the pre-antibiotic sample. In this study, we observed that the long-term effects of antibiotic treatment extend beyond transient microbial shifts; persistent changes in the plasmidome indicate a potential reservoir of antibiotic resistance genes, potentially accumulating and diversifying over time. Thus, monitoring plasmid populations post-antibiotic treatment is critical, as the colonization of novel resistance plasmids which may pose significant public health risks^115^.

To study the role of gut plasmidome in reserving antibiotic resistance genes within the microbiome, we extracted the AMR carrying sequences and assessed their prevalence through time. The *sul*2 gene commonly facilitates a diverse AMR profile, often co-transferring with other resistances^116^. This particular resistance gene, conferring sulfonamide resistance, is extensively studied in this genetic context (plasmid) and has been reported in clinical *E. coli* isolates at rates as high as >80% ^117–120^. Our findings, which indicate the flourishing of a specific plasmid subset during treatment, warrant further investigation into their evolution and dynamics. Another intriguing discovery was the increased prevalence of specific plasmid-encoded resistance genes (*cfxA3, cfxA6, erm*G, *met*A, and *msr*D) post-treatment. Notably, the *cfx*A6 gene, located both on chromosomes and plasmids, stood out with the significant increase in abundance after the antibiotic treatment. It is noteworthy that the patient underwent additional antibiotic therapy (cephalexin) for pharyngitis at month 10, which is likely to contribute to additional discrepancies within the gut microbiome. Despite this, our findings show stability of the microbial community between 180 and 365 days (Figure 3). When assessing the recovery of the plasmid population (180 and 365 days), the metagenomic analysis indicated significant diversification compared to day 7. The plasmid population remained relatively stable as it was highly similar at days 180 and 365. Further, new colonizing plasmids were rare during the last 6 months of the sampling period (Figure 3C).

Through the temporal analysis of the *E. coli* isolates and metagenomic data, an extensive but temporary domination of ESBL-*E. coli* (harboring both AmpC mutations and *bla*CTX-M-15) was observed during antibiotic therapy. This was further supported by repeated isolation of the same sequence type (ST648) before and during the treatment. However, after 180 and 365 days follow-up periods, the microbial composition and the plasmid population were partially restored and the ESBL-*E. coli* (*bla*CTX-M-15) was no longer detected. Instead, other *E. coli* sequence types of (ST69, ST73 and ST929), were seen to colonize the gut microbiome. All these isolates carried AmpC mutations and ST69 also plasmid-encoded betalactamases *bla*TEM-1B (isolate 6.2) and *bla*CMY-variant (isolates 6.2 and 12.1). Notably, these AMR genes were only found in follow-up isolates, and not detected via the metagenomic sequencing analysis at any timepoint. Hence, while metagenomics is an efficient tool for studying gut plasmidome, the scarce community members and their plasmids may remain undetected, especially when embedded in a diverse microbial community. This highlights the importance of conventional cultivation-based isolation strategies, which can serve as a complementary approach to capture microbial elements that might be overlooked or lost due to the inherent biases of metagenomic analysis^121^. Combining a metagenomic approach with culture-based methods could enhance the accuracy and completeness of the entire microbiome to better understand microbial dynamics in changing environments. Further, conventional metagenomic sequencing does not allow linking plasmids to their bacterial host cells as host species determination is often limited to the previously characterized plasmids and the available sequence databases. In addition to culture-based methods, recently established microbial single-cell methods, such as epicPCR^122^ and single-cell ddPCR^123^, and methods employing DNA cross-linking, such as HiC^124^, would provide more in-depth insights into identifying all the potential plasmid-carrying bacteria.

In conclusion, we observed a transient alteration in the gut plasmidome, AMR profile and their host microbial composition probably caused by the antibiotic treatment. A case study focusing on one patient has inherent limitations in establishing generalizable conclusions. Comparative analyses with control groups, such as those receiving placebo, are needed to confidently attribute disruptions and subsequent recovery of the microbiome to antibiotic treatment. Additionally, it needs to be acknowledged that appendicitis itself can significantly influence and cause the dysbiosis of the gut microbiome within a patient. This potentially confounds with the observed effects of antibiotic therapy. The gut plasmidome is a key source and mediator of the AMR spread among bacteria. Therefore, it is anticipated that the future research should assess plasmid dynamics during antibiotic therapies in broader clinical contexts to fully address the role of plasmid-host interplay in maintaining AMR gene pools within human microbiomes.

## Supplementary Materials

**Supplementary Table S1.** Antibiotic susceptibility and mobility testing of the *E. coli* isolates.

| Isolate | Resistances |  |  |  |  |
| --- | --- | --- | --- | --- | --- |
|  | Ampicillin | Mobility | Cephalothin | Mobility | Rifampicin |
| 0.1 | S | - | S | - | S |
| 0.2 | R | - | R | - | S |
| 1.1 | R | - | R | - | S |
| 1.2 | S | - | S | - | S |
| 3.1 | R | - | R | - | S |
| 3.2 | R | - | R | - | S |
| 7.1 | R | - | R | - | S |
| 7.2 | R | - | R | - | S |
| 6.1 | S | - | S | - | S |
| 6.2 | R | - | R | - | S |
| 12.1 | R | - | R | - | S |
| 12.2 | R | - | R | - | S |

**Supplementary Table S2.** Plasmid characteristics of *E. coli* isolates. Metagenome indicates if the plasmid was found from metagenome data. Host isolate indicates the isolates found to carry the plasmid. PTU = plasmid taxonomic unit.

| Plasmid | Metagenome | Host isolate | Replicon | PTU | ESBL | AMR |
| --- | --- | --- | --- | --- | --- | --- |
| 1 | False | 6.2,12.1 | IncB/O/K/Z | PTU-B/O/K/Z | *bla*CMY | aph(6)-Id ,<br>aph(3'')-Ib,<br>sul2, |
| 2 | False | 6.2,12.1 | IncX4 | PTU-X4 | - | - |
| 3 | False | 6.2,12.1 | Col8282 | PTU-E10 | - | - |
| 4 | False | 6.2,12.1 | - | PTU-E1 | - | - |
| 5 | True | 0.2, 1.1, 3.1,<br>3.2, 7.1, 7.2,<br>12.2 | IncFII | PTU-FE | - | - |
| 6 | True | 0.2,<br>1.1,3.1,3.2,7.<br>1,7.2 | IncI2 | PTU-I2 | - | - |
| 7 | True | 0.2, 1.1, 3.1,<br>3.2, 7.1, 7.2,<br>12.1 | Col156 | PTU-E7 | - | - |
| 8 | True | 0.2,<br>1.1,3.1,3.2,7.<br>1,7.2 | Col(BS512) | PTU-E47 | - | - |
| 9 | True | 12.2 | IncFIB(AP001918) | PTU-FE | - | - |
| 10 | False | 12.2 | IncFII(29) | NA | - | - |
| 11 | False | 6.2 | IncFII | PTU-FE | - | - |

**Supplementary Figure S1.**
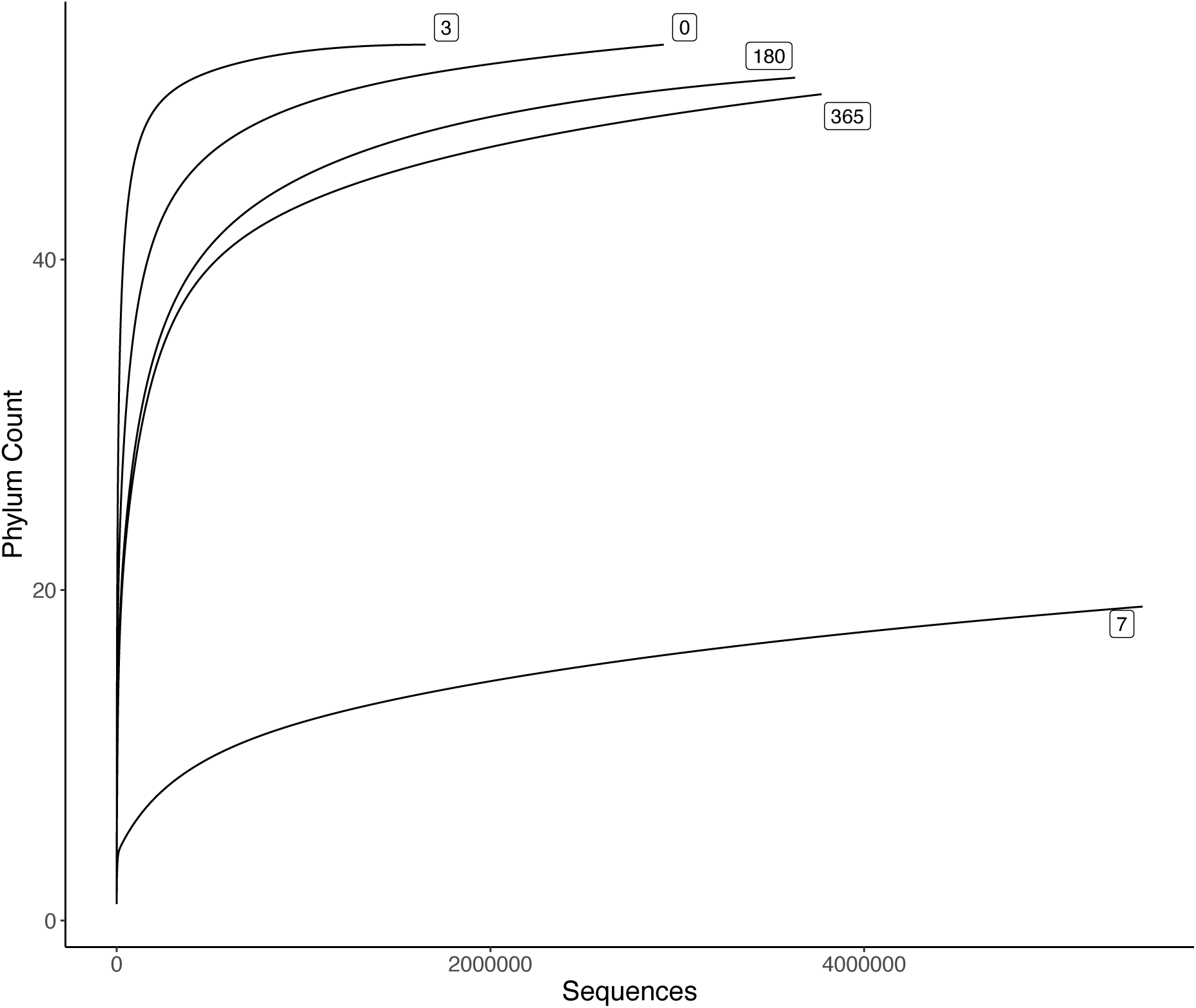
Rarefraction curve of the microbial community during a 7-day antibiotic treatment (sampled at days 0, 3, 7, 180 and 365).

**Supplementary Figure S2.**
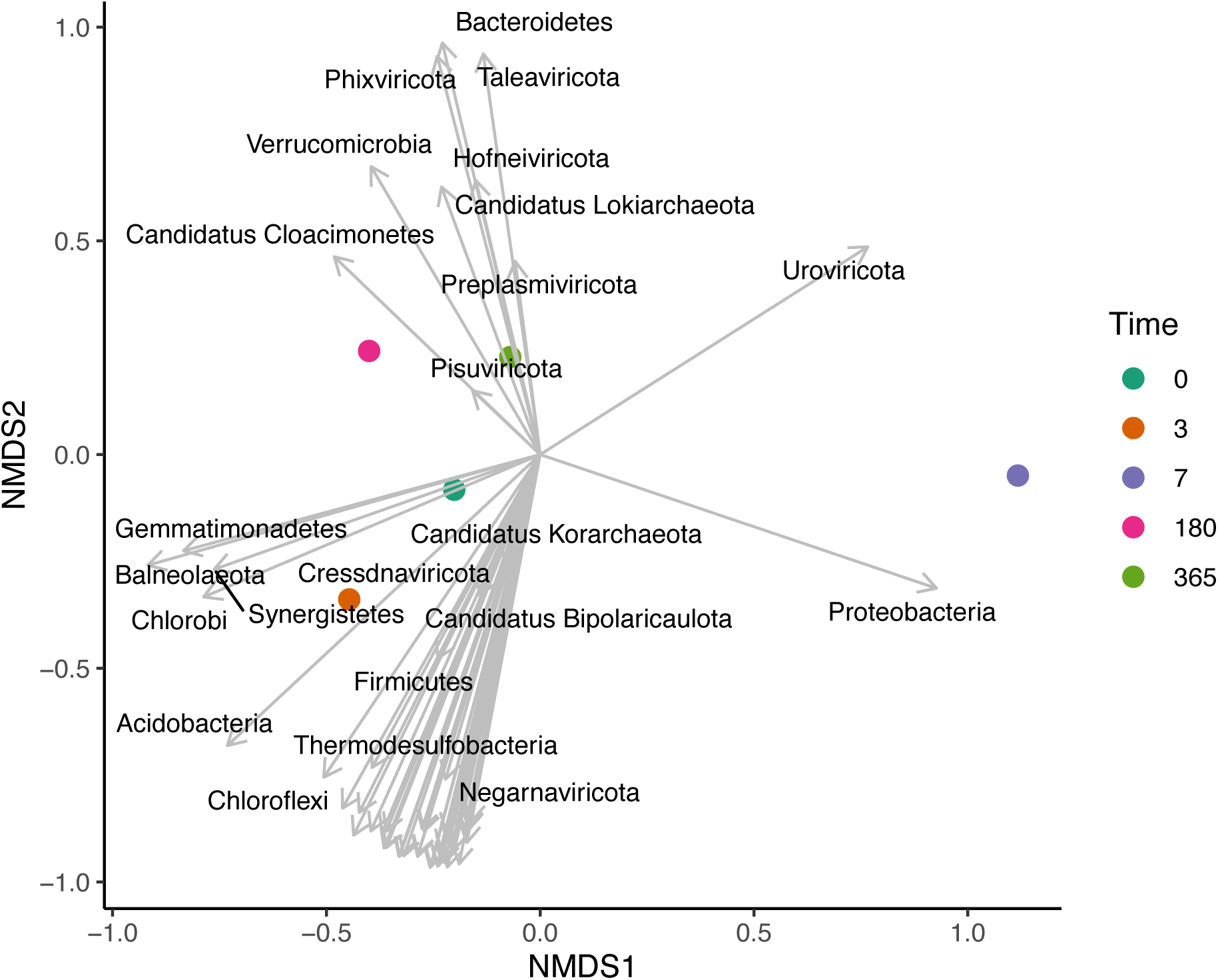
Microbial taxa explaining the differences in community structure during a 7-day antibiotic treatment (sampled at days 0, 3, 7, 180 and 365).

## Acknowledgements

The authors would like to acknowledge Anna Musku for assistance in the laboratory work. The study was funded by Research Council of Finland (grants #322204 and #354982 to RP, #346772 for L-RS ). S.M. is supported by funding from the Biotechnology and Biological Sciences Research Council [BB/X009793/1].

## Author contributions

Original idea: RP, AJH, PS

Study design: RP, SM, IJ, TK, AP, SV, AJH, PS

Data collection: IJ, RP, AP, SV, AJH, PS, MJ

Data analysis: IJ, RP, SM, JR, EW, SVH

Writing: IJ, RP, SM, TK, AP, SV, AJH, PS, MJ, SM, EW, SVH, LRS, JR

IJ = Ilmur Jonsdottir, RP = Reetta Penttinen, MJ = Matti Jalasvuori, LRS = Lotta-Riina Sundberg, TK = Teemu Kallonen, AP = Annaleena Pajander, SV = Sanja Vanhatalo, AJH = Antti J. Hakanen, PS = Paulina Salminen, SM = Sean Meaden, EW = Edze Westra, SVH = Stineke van Houte, JR = Janne Ravantti

